## Supplementary text for "Lineage priming and cell type proportioning depends on the interplay between stochastic and deterministic factors"

### Expectation of the stochastic-deterministic model at its limits

In the main text we note that the stochastic-deterministic model can produce anywhere from the step-like function expected under the deterministic model (eqn. 1 and Figure 1A) and the constant proportioning expected for a purely stochastic model (equation 2 and Figure 1B). The case of the purely deterministic model can be derived by taking the limit where variation from noisy gene expression vanishes, which occurs as the variance in gene expression goes to zero (i.e., as $\sigma \to 0$):

$$P_{t(\text{deterministic})} = \frac{1}{2}\left(1 + \frac{\sqrt{(C_0 - R - \beta t)^2}}{C_0 - R - \beta t}\right) \quad \text{(S1)}$$

This expression takes on the value of 1 during the period of the cell cycle where $t < (C_0 - R)/\beta$ and a value of 0 otherwise. The case of the purely stochastic model can be derived by taking the limit where the cell-cycle dependent component vanishes (i.e., as $\beta \to 0$), which yields the expression given by equation (2), where stalk propensity depends on the properties of the stochastically expressed genes ($\mu$ and $s$) relative to the threshold that results in stalk fate ($R$).

### Comparison of model fit to the exponential-decay model of Gruenheit et al.

*Model fitting methods*

In the main text we evaluate the fit of the stochastic-deterministic model to the stalk propensity data from (Gruenheit et al., 2018), and compare it to their exponential-decay model. Here we provide further information on the exponential-decay model and the methods used to compare the fit of the two models.

Like the stochastic-deterministic model, the exponential-decay model is a two-parameter model that predicts stalk propensity through the cell cycle. It treats stalk propensity as a deterministic property that changes through the cell cycle according to the equation $\tilde{P}_t = \chi \exp(-\lambda t)$, where $\chi$ represents the starting stalk propensity (just after the end of mitosis), $\lambda$ the exponential rate of decay in stalk propensity, and, $t$ the time since the end of the last mitosis (with the tilde being used to differentiate the expectation for this model from the expectation from the stochastic-deterministic model). Although both models have two parameters that capture similar properties, they are not equivalent to each other. The stochastic-deterministic model defines a starting stalk propensity as a Z-score ($C_0^*$) that translates into the proportion of cells from a stochastically variable (Gaussian) distribution that will become stalk, whereas the exponential-decay model defines a fixed starting propensity ($\chi$). Both models have decay terms, with the decay term in the exponential-decay model ($\lambda$) giving the exponential rate of decrease in stalk propensity, while the decay term in the stochastic-deterministic model ($\beta$) represents the decay in the standardized mean CCAF level.

The exponential-decay model was fitted to the data following the same method as that described in the text for the fitting of the stochastic-deterministic model. Briefly, the model was fitted using the 'NonlinearModelFit' function in Wolfram Mathematica version 14, which uses the Levenberg–Marquardt algorithm for least-squares curve fitting. This generated estimates for the two parameters, $\chi$ and $\lambda$ and for the Akaike Information Criterion (AIC) of the fitted model. We used the error and corrected total sums of squares to calculate the $R$-squared (since it is calculated incorrectly by default in the NonlinearModelFit function).

We evaluated the relative support for each model by comparing the Akaike weights (Burnham et al., 2002; Wagenmakers and Farrell, 2004), which reflect the relative likelihood of the models. To further evaluate the relative fit of the models, we used the parameter estimates to generate a set of predicted values for each model and treated these as independent variables in a single linear model where the observed (adjusted) stalk propensity values were the dependent variable, which essentially allows the two models to directly compete for the fit to the data.

*Model fitting results*

As expected based on the findings of Gruenheit et al. (2018), the exponential-decay model provides a good fit to the data (adjusted $R$-squared = 0.88, $AIC = -47.19$), yielding an estimated rate of decay of $\lambda = 0.351$ and a starting stalk propensity of $\chi = 0.74$ (Supplementary Figure 1.A.i), which would correspond to an overall steady-state stalk propensity of 0.35 (where all times since the end of the previous mitosis are sampled equally). As noted in the main text, although the model provides a good fit to the data, it does not offer a mechanism to translate CCAF levels into quantitative variation in stalk propensity. Hence, it provides a description of the pattern of change in stalk propensity through the cell cycle (as being approximately an exponential decline) rather than a mechanistic model for the process that produces the predicted value of stalk propensity.

The ratio of the Akaike weights of the stochastic-deterministic model compared to the exponential-decay model has the value 98.35, which implies that the stochastic-deterministic model is 98.35 times more likely (in terms of Kullback–Leibler discrepancy). This result can also be interpreted as a probability of 0.990 that the stochastic-deterministic model is the better fitting model (meaning we can reject the exponential-decay model as the better fitting model with $p = 0.010$). This conclusion is further supported by the results of the linear model that competed the two sets of predictions, which indicated that the partial fit to the predicted values from the stochastic-deterministic model is very good ($F_{1,20}$ = 9.83, $p$ = 0.0052) while the fit to the predicted values from the exponential-decay model is not significant ($F_{1,20}$ = 0.48, $p$ = 0.49).

**Evaluating the assumption of normally distributed noise**

To evaluate the assumption of normality for the distribution of noise in the stochastic-deterministic model we used an approach that models noise based on the gamma distribution, which is a family of continuous distributions that vary in the values of two parameters (in our implementation, these are a shape parameter $k$ and a scale parameter $\theta$). By varying the two parameters, the gamma distribution provides probability density functions that goes from a pattern of exponential decay to a pattern that approximates normality. The gamma distribution does not have a fixed mean and variance, but the shape parameters together dictate these two properties of the distribution, where the mean is given by $k\theta$ and variance by $k\theta^2$. Using these properties, we modelled decay in CCAF by adapting the gamma distribution to a form that is directly analogous to the approach implemented in the stochastic-deterministic model:

$$P_t = \frac{\theta^{-k}}{\Gamma(x)} \int_R^{\infty} e^{-\frac{x}{\theta}} x^{k-1}\, dx = \frac{\Gamma\left(k, \frac{C_0 + \beta t}{\theta}\right)}{\Gamma(k)} \qquad \text{(S2)}$$

where $\Gamma(x)$ is the Euler gamma function, and $\Gamma(x, y)$ is the incomplete gamma function (which is integrated from $y$ to ∞).

Following the methods used to fit the exponential-decay and stochastic-deterministic models, we again used the 'NonlinearModelFit' function in Wolfram Mathematica version 14 to fit the gamma-distribution model (eqn. S2) to our data, except the model includes a total of four parameters ($C_0$, $\beta$, $k$ and $\theta$) instead two parameters like the other two models. This not only allows us to evaluate whether the best fit function was consistent with the assumption of normality, but also to further evaluate the Gaussian noise model against the exponential-decay model (since the best fit gamma distribution could approximate either distribution). We find that the best fit model ($C_0$ = 1.227, $\beta$ = 0.074, $k$ = 56.9, and $\theta$ = 0.023) has a good fit to the data (adjusted $R$-squared = 0.91, $AIC$ = –51.60; see Supplementary Figure 1A.ii), with a gamma distribution component corresponding to a distribution of noise that is approximately normal (cf. Supplementary Figure 1.B.i and 1B.ii). Hence, the additional two parameters for the gamma distribution yield a similar fit to the simpler stochastic-deterministic model that assumes Gaussian noise, meaning that gamma-distribution model will necessarily be the poorer fitting model given it has two additional parameters (so we do not include any formal model comparisons). This conclusion is further supported by the ratio of the Akaike weights of the model with Gaussian noise compared to the model with gamma distributed noise, with has the value 10.85, meaning that, despite having two fewer parameters, the Gaussian noise model is 10.85 times more likely (corresponding to a probability of 0.92).

**Evaluating the assumption of linear decay in CCAF**

The simple version of the stochastic-deterministic model presented in the main text assumes that CCAF shows a linear decay through the cell cycle. What this means in terms of the model properties is that the mean of the distribution of stalk-inducing factors changes linearly through the cell cycle (since the level of CCAF determines the mean level of stalk inducing factors, while CCIF adds variability around this mean). Importantly, this assumption does not necessarily imply that there is a linear change in some factor(s) at the molecular level (since the number of molecules of a signal could decay exponentially, but the effect of the signal could decay linearly if there is a non-linear relationship between signal levels and their effect, and vice versa). For many biological processes, we might expect factors to show non-linear decay, especially exponential decay (e.g., if concentrations showed a constant half-life). Therefore, to evaluate whether a model with non-linear decay provides a better fit to the data, we replaced the linear decay function in equation (1) with an exponential decay function. Initial attempts to fit a simple two parameter decay process (such as the exponential decay equation fitted in Gruenheit et al., which we could write as $A_0 e^{-\delta t}$) made it clear that there was no parameter space that would produce the pattern of change in stalk proportioning observed through the cell cycle. This issue is due to the fact that the simple exponential decay equation ties together the 'starting value' (which would be $A_0$ when $t = 0$) and the effective rate of decay (e.g., the derivative of the decay function with respect to time, would be $-\delta A_0 e^{-\delta t}$, so the linear component depends on the decay rate $\delta$ scaled to the starting value $A_0$). Therefore, we used a more flexible exponential decay function that allows for separation between the starting value and the rate of decay:

$$A_t = A_0 - A^* e^{-\delta t} \qquad \text{(S3)}$$

The structure of equation (S3) is superficially similar to the linear decay equation (eqn. 2). However, while $A_0$ is the starting level of CCAF in equation (1) (i.e. the level of CCAF that determines stalk proportioning at the start of the cell cycle), in equation (S3) the starting level (where $t = 0$) would be $A_0 - A^*$. The level of CCAF is eroded by an exponential process captured by the second term on the

RHS, which has two parameters, $A^*$, which represents the size of the CCAF pool that decays, and $\delta$, which gives the rate of exponential decay.

Following the same logic outlined in the main text for the case of linear decay in CCAF, we assume stalk proportion depends on the proportion of cells experiencing a value of CCAF + CCIF that is above a threshold value. Replacing linear decay with the exponential decay process in equation (S3) gives $C_t = \mu + A_0 - A^* e^{-\delta t} = C_0 - A^* e^{-\delta t}$. As in the linear decay model, we can rescale the value of $C_0$ relative to $R$, though the interpretation of the resulting parameter $C_0^*$ is different in this case since it does not, on its own, capture the baseline value that represents the expectation for stalk propensity at the start of the cell cycle. Instead, while $C_0^*$ gives the baseline value in the linear decay model (where $C_0^* = R - (\mu + A_0) = R - C_0$), in the exponential decay model the baseline value (i.e., the value that determines stalk fate at the start of the cell cycle) is given by $C_0^* - A^*$. Using this value of $C_0^*$ in place of the one for the linear model to derive the stalk propensity gives (cf. eqn. 3):

$$P_t = \frac{1}{2}\left(1 - e^{\delta t}\sqrt{e^{-2\delta t}}\right) \operatorname{erf}\left[\frac{\sqrt{e^{-2\delta t}}\left(A^* - C_0^* e^{\delta t}\right)}{\sqrt{2}}\right] \quad \text{(S4)}$$

Because of the more complex structure of equation (S4), this model was fitted using the 'NMinimize' method in the 'NonlinearModelFit' function in Wolfram Mathematica version 14, which forces a global search for the best fit parameters and helps avoid local minima (note that using this method for all other models fitted in our study has no impact on the estimates). The exponential decay model shows a slightly worse fit than the linear decay model (adjusted *R*-squared = 0.920 and AIC = −56.37 for the linear decay model, while adjusted *R*-squared = 0.916 and AIC = −54.24 for the exponential decay model), but given that the fits are almost identical, and the linear decay model shows a slightly better fit, the linear decay model is necessarily the more likely model (with the ratio of the Akaike weights being 2.90). Moreover, the two models produce very similar estimates for the starting level of CCAF (which is an estimate of $C_0^*$ in the linear decay model and $C_0^* - A^*$ in the exponential decay model), with values of 0.57 for the linear model and 0.56 in the exponential model. They also produce similar estimates for the linear rate of decay (which is given by the term $\beta$ in the linear model, and the first derivative of the solution with respect to $t$ in the to the exponential decay model), with values of 0.41 for the linear model and 0.39 for the exponential model. Importantly, the estimates from the exponential decay model produce a pattern of decay in CCAF that is almost perfectly linear (see Supplementary Figure 1.C.i), which reflects the fact that the quadratic change in CCAF (given by the second derivative of the solution with respect to $t$ in the to the exponential decay model), which has the value 0.01, is tiny compared to the linear change (as are all higher order relationships). Hence, the fact that the parameter estimates from the best fit exponential decay model produce an almost perfectly linear change in CCAF through the cell cycle, while showing a model fit that is slightly worse than the linear decay model despite having an additional parameter, provides strong support in favour of the linear decay model over an exponential decay model.

To further support this conclusion, we fitted two other generic models for decay of CCAF in the stochastic-deterministic model, a quadratic model of change in CCAF (i.e., where CCAF can change as a function of $t$ and $t^2$) and a cubic model of change in CCAF (i.e., where CCAF can change as a function of $t$, $t^2$ and $t^4$). The quadratic model shows a similar fit to the stochastic-deterministic model with exponential decay in CCAF (adjusted *R*-squared = 0.916 and AIC = −54.25 for the quadratic decay model, cf. above). The reason the fits of the quadratic and exponential decay models are so similar is because the best fit parameters for both models effectively produce linear decay (cf. Supplementary Figures 1.i and 1.ii), which is reflected in the fact that the estimated quadratic term is near zero (with a value of 0.007). Like the stochastic-deterministic model with exponential decay, the model with quadratic decay is less likely than the model with linear decay because it has an additional parameter (with the ratio of the Akaike weights being 2.88). The cubic model shows a similar fit to the stochastic-deterministic model with linear decay (adjusted *R*-squared = 0.921 and AIC = −54.64), but again, because it requires two additional parameters it is still the less likely model (with the ratio of the Akaike weights being 2.37). Hence, although some degree of non-linearity appears in the best fit parameters for the cubic model (see Supplementary Figure 3.iii), it does not improve the fit of the model, and hence the most likely model is one with linear decay.

**Model fitting with the outlier value included**

To confirm that removal of the outlier stalk propensity value measured for *gefE*⁻ under G− at hour 5 did not alter our results we fitted the exponential-decay and stochastic-deterministic models to the full dataset. The exponential-decay model shows the same fit in terms of adjusted *R*-squared as the model with the outlier removed (adjusted *R*-squared = 0.88, *AIC* = –50.61), yielding a similar estimated rate of decay ($\lambda$ = 0.345 compared to 0.351 for the model with the outlier removed) and an identical starting stalk propensity ($\chi$ = 0.74 for both datasets). Likewise, the stochastic-deterministic model shows approximately the same fit in terms of adjusted *R*-squared as the model with the outlier removed (adjusted *R*-squared = 0.91, *AIC* = –57.47), yielding almost identical estimates for starting sensitivity ($C_0^*$ = 0.557 compared to 0.574 for the model with the outlier removed) and rate of decay in sensitivity ($\beta$ = 0.397 compared to 0.409 for the model with the outlier removed). The ratio of the Akaike weights of the models with the outlier included is smaller than for the analysis with the outlier removed (30.91 vs 93.46), but still corresponds to a probability of 0.969 that the stochastic-deterministic model is the better fitting model (meaning we can reject the exponential-decay model as the better fitting model with *p* = 0.031).

**Influence of stochastic variation on sensitivity to cell-cycle perturbations**

To consider how the presence of stochastic variation from noisily expressed genes can buffer against the impact of cell-cycle perturbations on stalk propensity we developed a simple model for biased sampling across the cell cycle. In the absence of bias, we assume that cells are sampled from a continuous uniform distribution from times zero to one, $U[0,1]$ (so the probability density is the same for all $t$, i.e., $f(t) = 1$ such that $\int_0^1 f(t)\,dt = 1$, meaning that time in the cell cycle is measured as the proportion of the cycle completed (e.g., if $t$ = ½, then cells would be halfway through the cell cycle, since $\int_0^{0.5} f(t)\,dt = 0.5$. The expected stalk propensity ($P_t$) at time $t$ is given by equation (3), so the expected propensity of a population of cells sampled uniformly across the cell cycle (denoted $P_u$) is $P_u = \int_0^1 P_t f(t)\,dt$.

Because our primary interest is on the impact of cell-cycle biases, and there is an unlimited range of possible ways for non-random sampling of cells from the cell cycle, we use a simple approach to capture non-random sampling across the cell cycle. We assume that bias is linear and defined by the equation $Q_t = 2\delta t - \delta$, where $Q_t$ gives the relative change in probability density caused by sampling bias, and $\delta$ measures the degree of sampling bias and ranges from –1 (bias towards the first half of the cell cycle) and +1 (bias towards the second half of the cell cycle). The degree of bias is translated into a bias in probability density function for sampling from the cell cycle as: $N(t) = f(t) + Q_t f(t)$ (see Supplementary Figure 12 for an illustration of the resulting biased probability density functions). This simple linear modification of the uniform probability density function retains the property that $\int_0^1 N(t)\,dt = 1$ (and hence still represents a probability density function) because it achieves a symmetrically re-distribution of probability density across the range of $t$ from 0 to 1. This makes the expected propensity of a population of cells sampled non-uniformly across the cell cycle (denoted $P_d$): $P_d = \int_0^1 P_t N(t)\,dt$.

There are a number of ways to consider the impact of this bias on the distribution of cells. For simplicity we consider the proportion of cells that were sampled from the first or second half of the cell cycle, with the proportion of cells in the first half of the cell cycle being $(2 - Q_t)/4$, meaning that, at the maximal degree of bias towards the first half of the cell cycle ($Q_t = -1$), ¾ of cells would be in the first half of the cell cycle (which we denote in Supplementary Figure 13 as –¾). We measured the impact of non-random sampling as the relative change in stalk proportioning: $(P_d - P_u)/P_u$. To consider how stochastic variation impacts the relative change in stalk proportioning we varied the value of $\sigma$, which gives the standard deviation of the distribution of noisy expression (see Supplementary Figure 13).

Burnham, K.P., Anderson, D.R., and Burnham, K.P. (2002). Model selection and multimodel inference : a practical information-theoretic approach, 2nd edn (New York: Springer).
Gruenheit, N., Parkinson, K., Brimson, C.A., Kuwana, S., Johnson, E.J., Nagayama, K., Llewellyn, J., Salvidge, W.M., Stewart, B., Keller, T.*, et al.* (2018). Cell Cycle Heterogeneity Can Generate Robust Cell Type Proportioning. Developmental Cell *47*, 494–508.e494-494–508.e494.
Wagenmakers, E.J., and Farrell, S. (2004). AIC model selection using Akaike weights. Psychon Bull Rev *11*, 192-196.
