## Supplementary figures for "Lineage priming and cell type proportioning depends on the interplay between stochastic and deterministic factors"

### Supplementary Figure 1

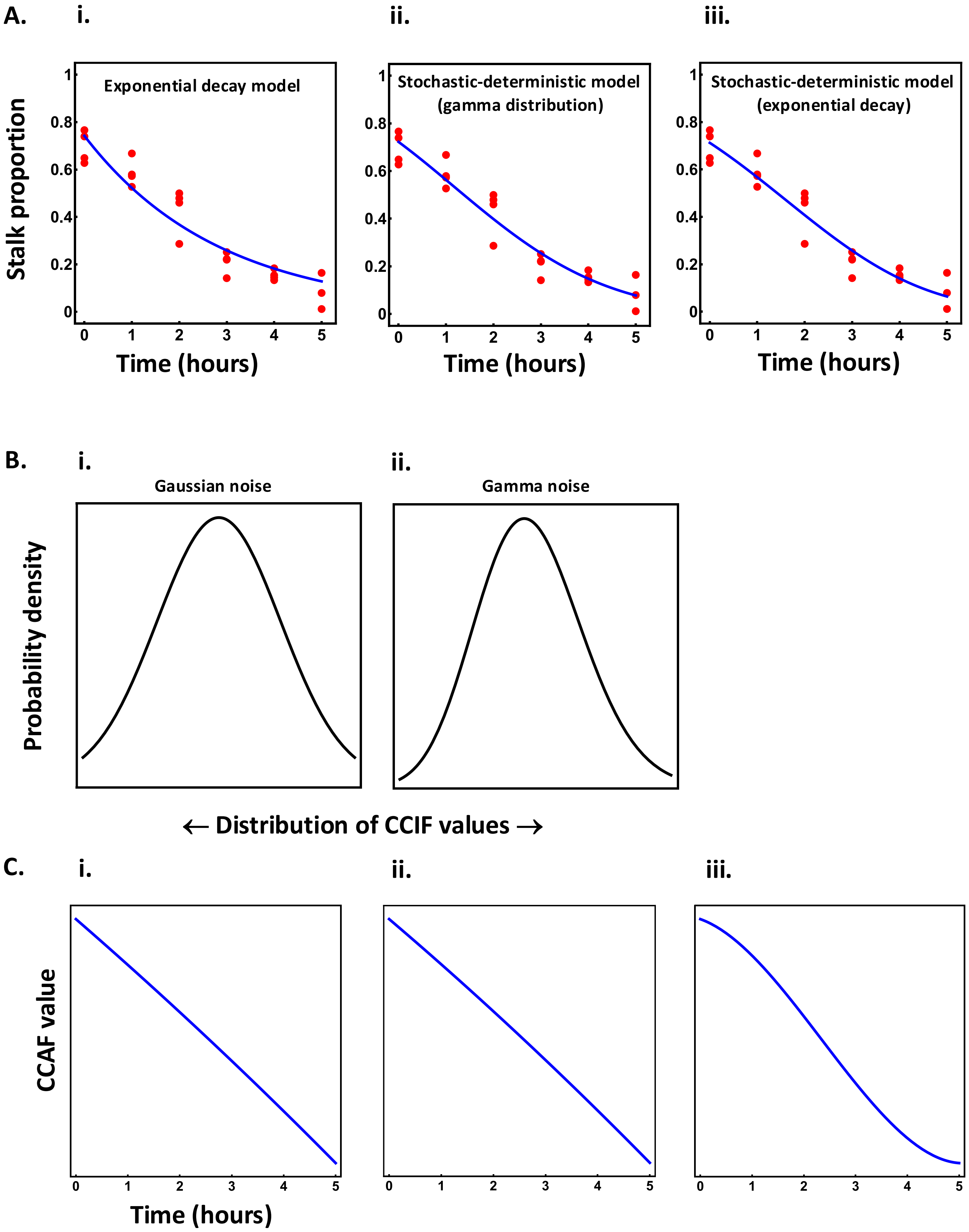

### Supplementary figure 2

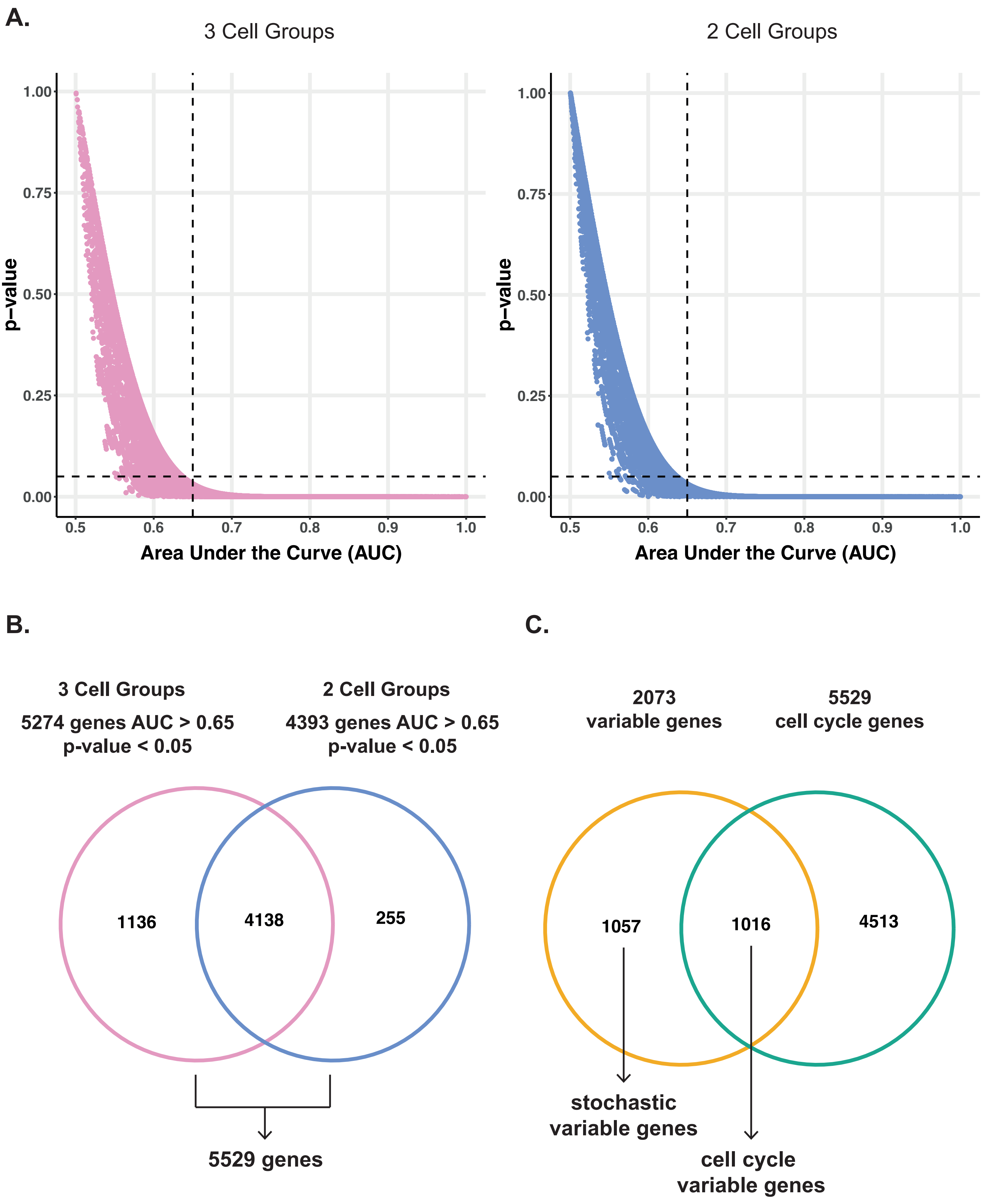

**Supplementary figure 3**

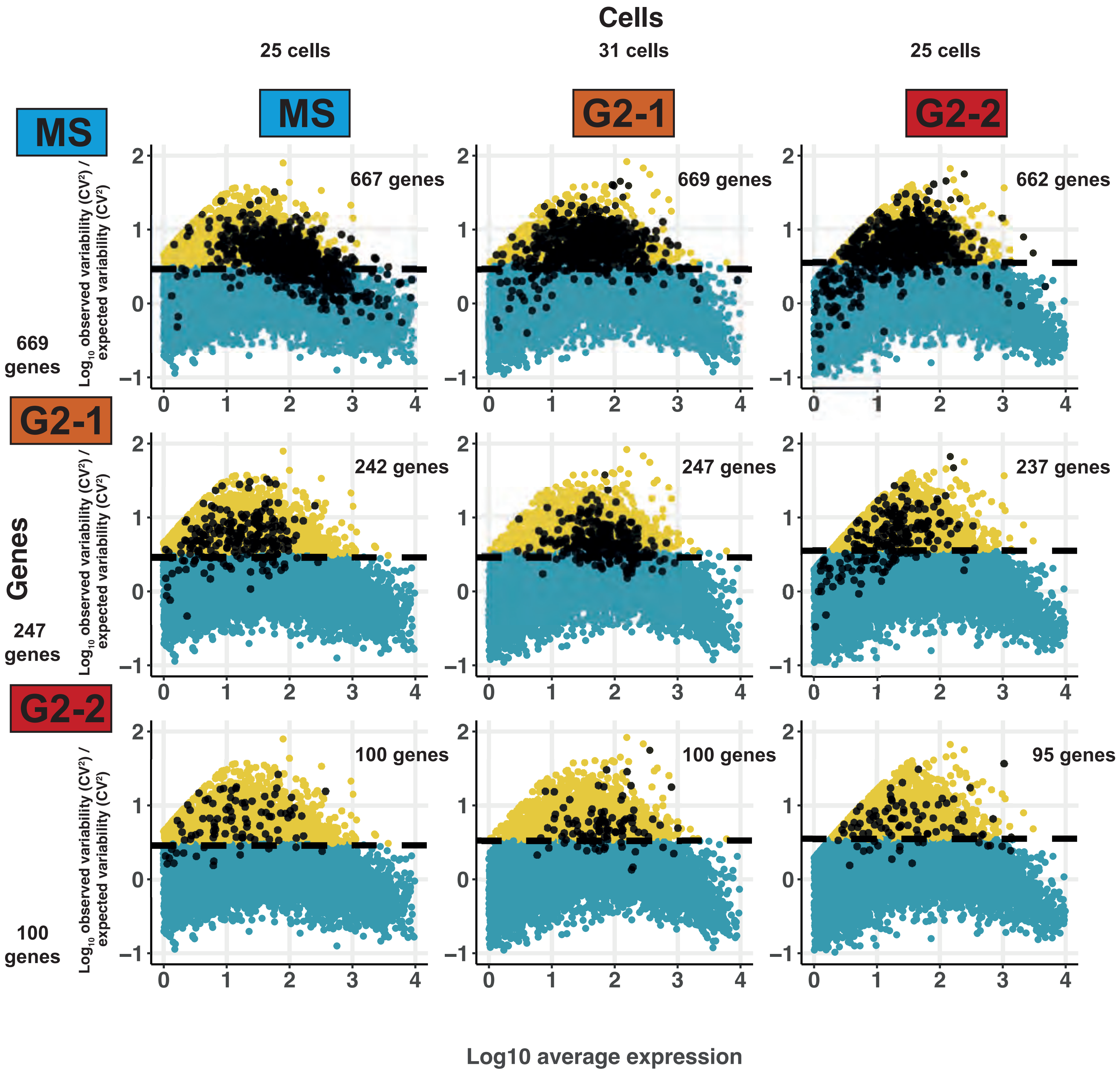

#### Supplementary figure 4

**A.**

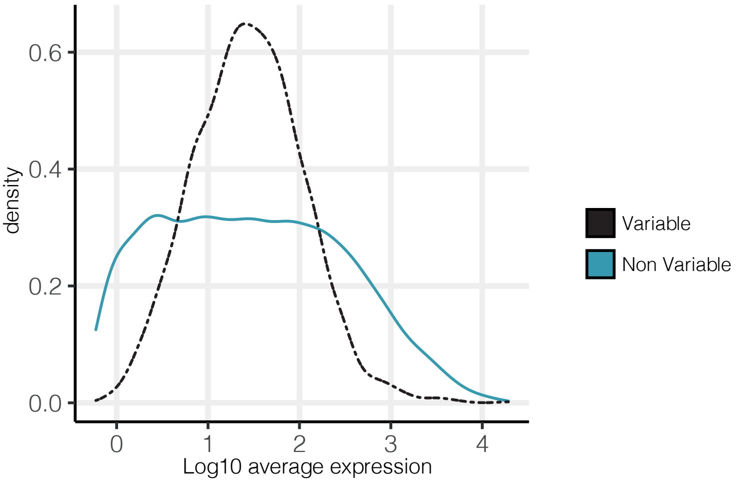


**B.**

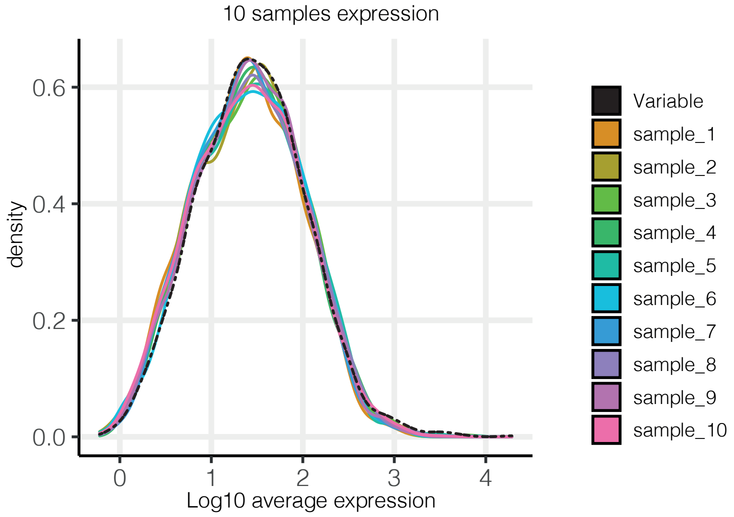


**C.**

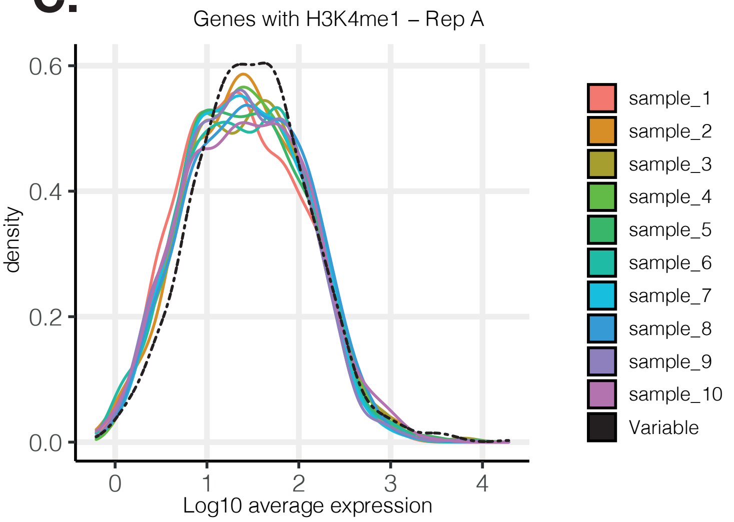


**D.**

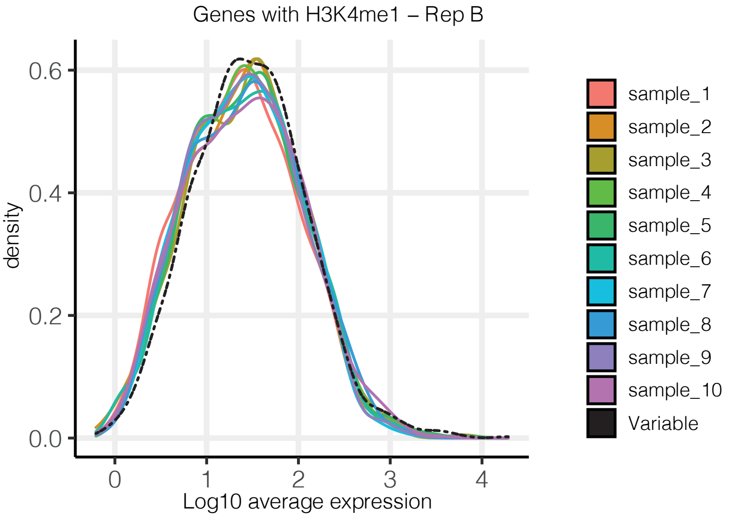


**E.**

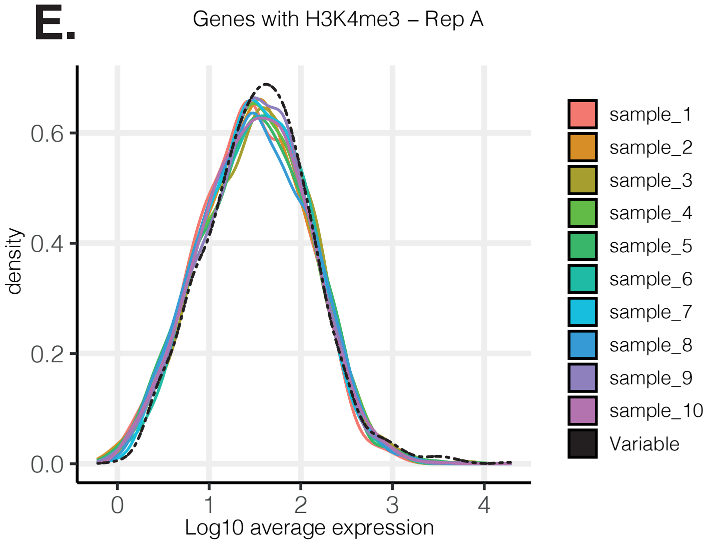


**F.**

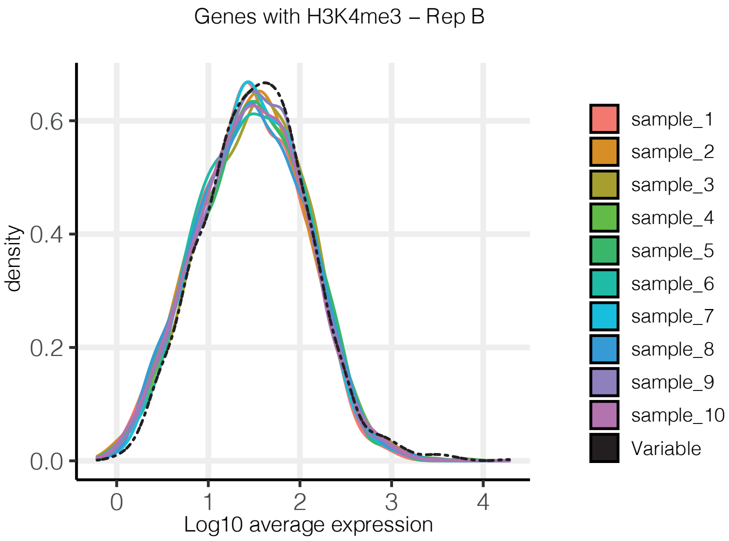


**G.**

| | Replicate_A | Replicate_B |
|---|---|---|
| *Ten Samples of Non Variable Genes with H3K4me1– Averaged* | 780.9 | 1005.4 |
| *Variable Genes with H3K4me1* | 1121 | 1349 |

**H.**

| | Replicate_A | Replicate_B |
|---|---|---|
| *Ten Samples of Non Variable Genes with H3K4me3– Averaged* | 1370.5 | 1630.1 |
| *Variable Genes with H3K4me3* | 1233 | 1505 |

Supplementary figure 5

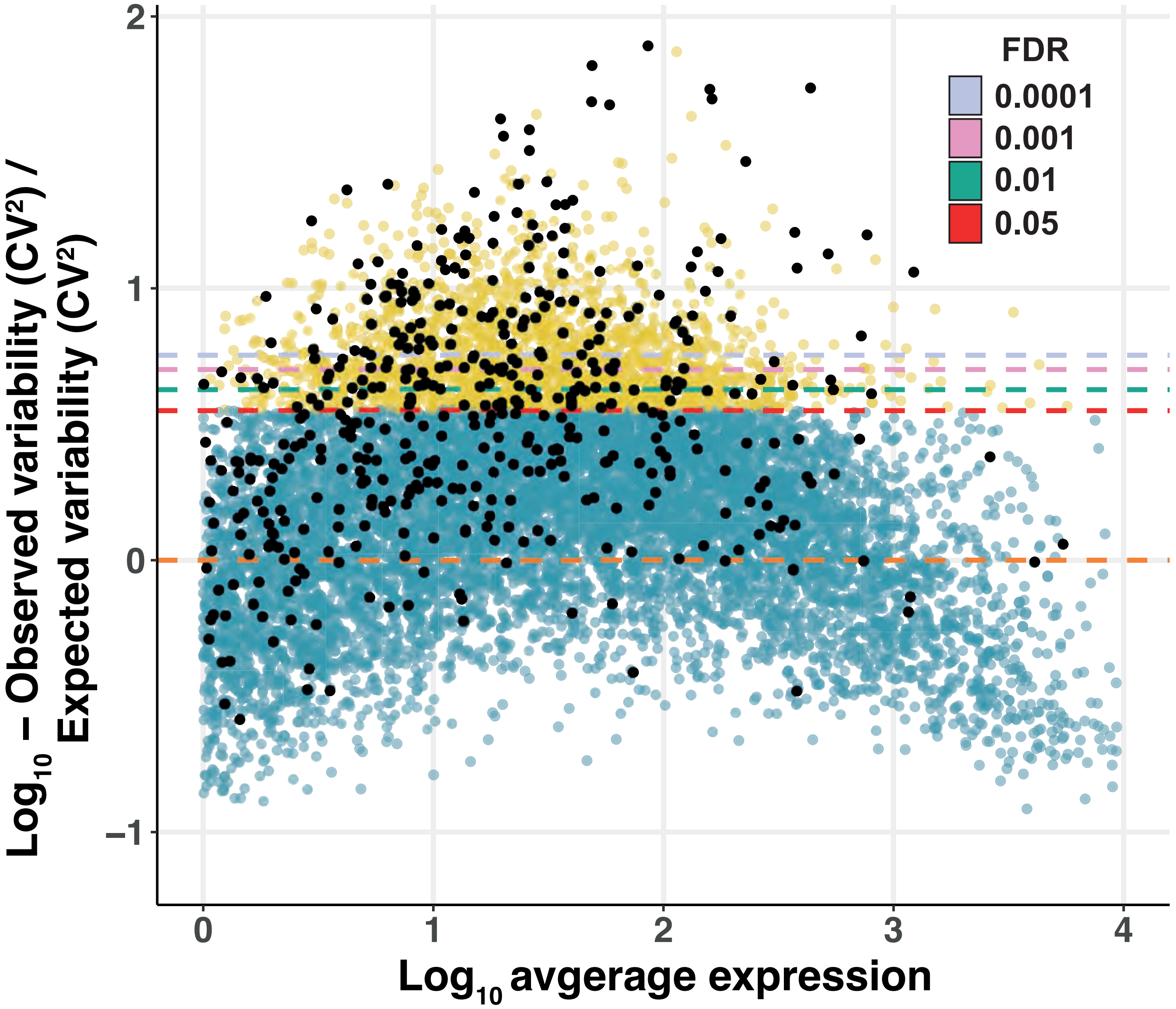

#### Supplementary figure 6

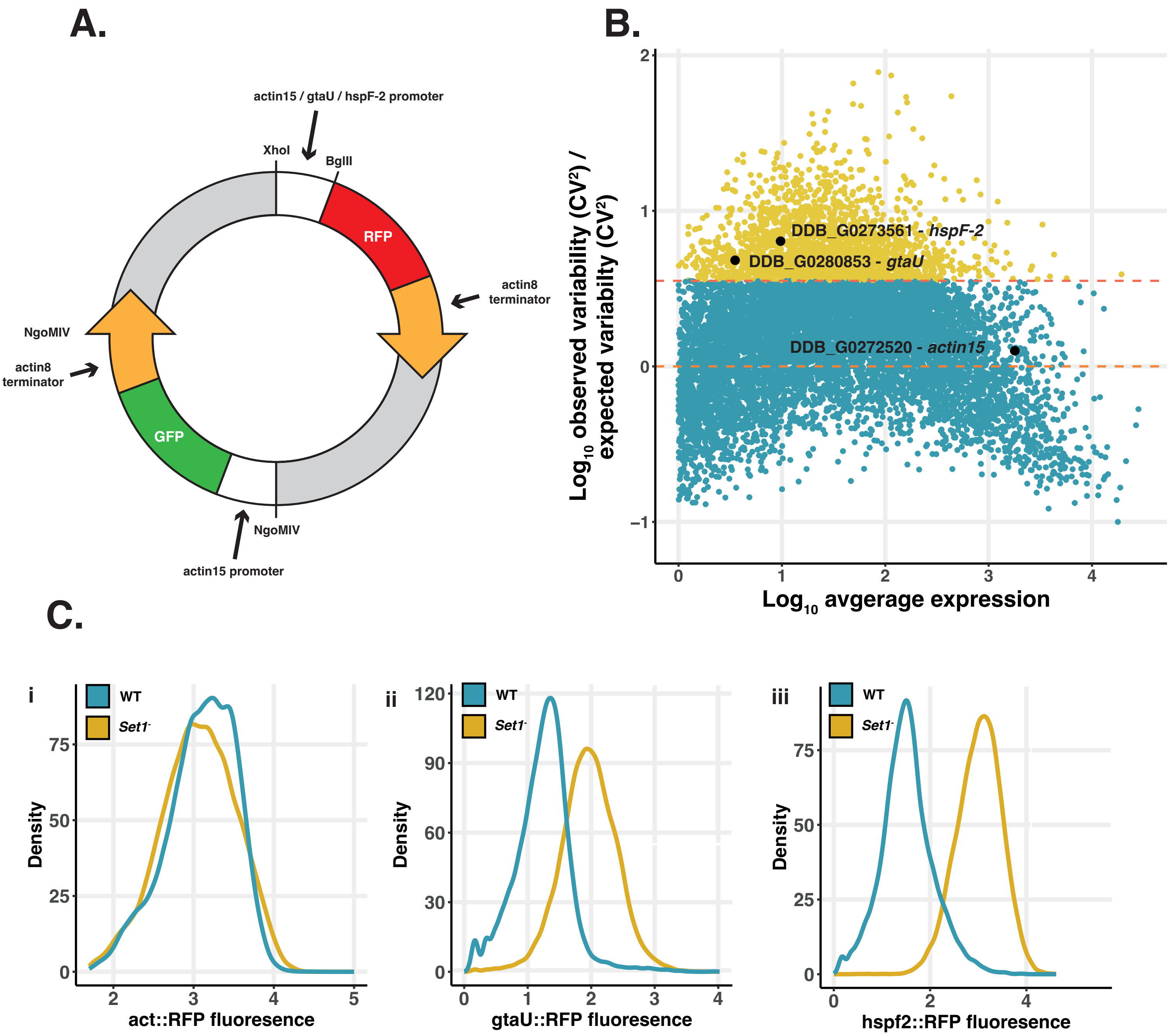

### Supplementary figure 7

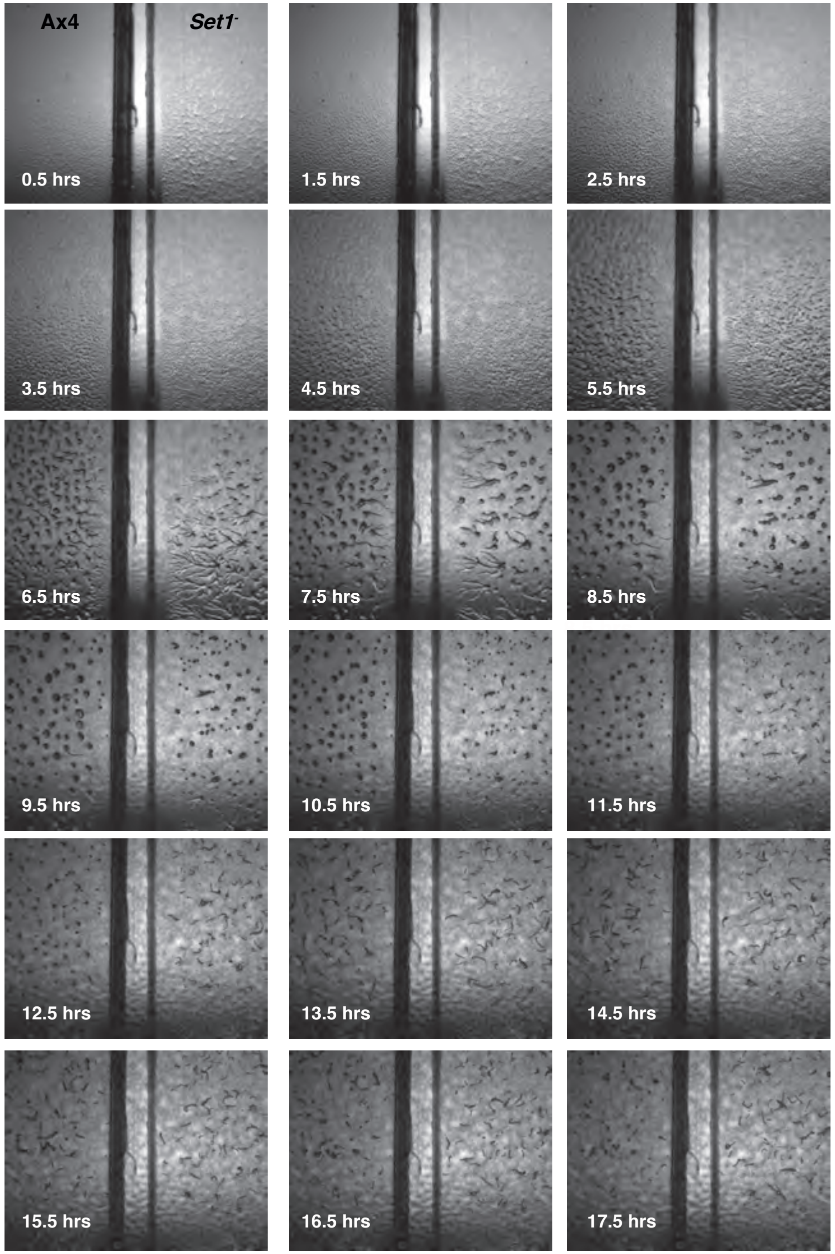


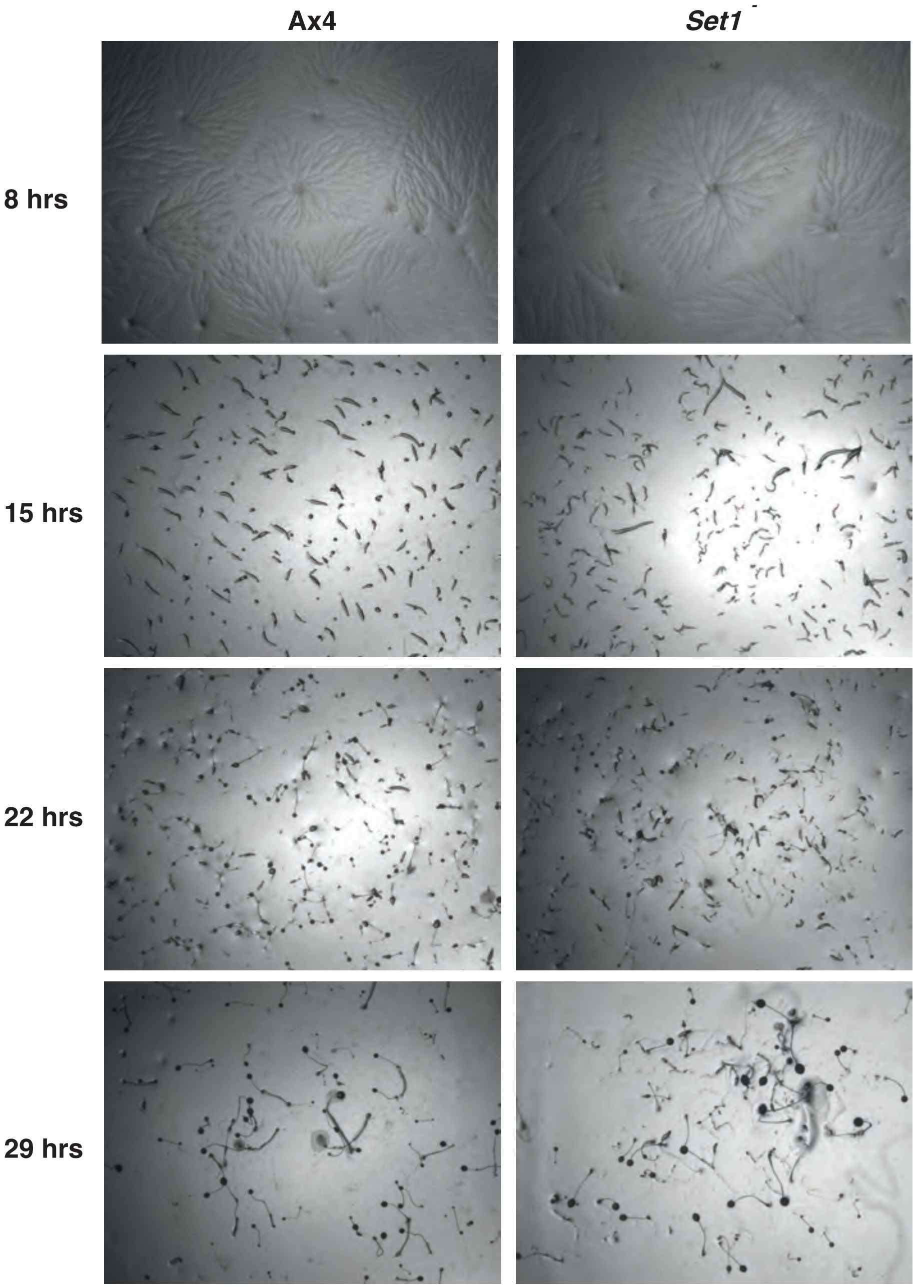

### Supplementary figure 8

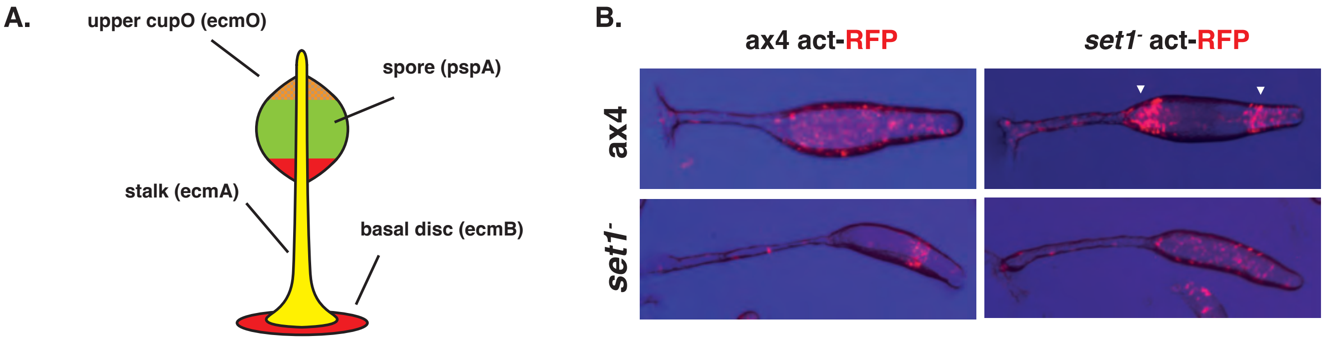

### Supplementary figure 9

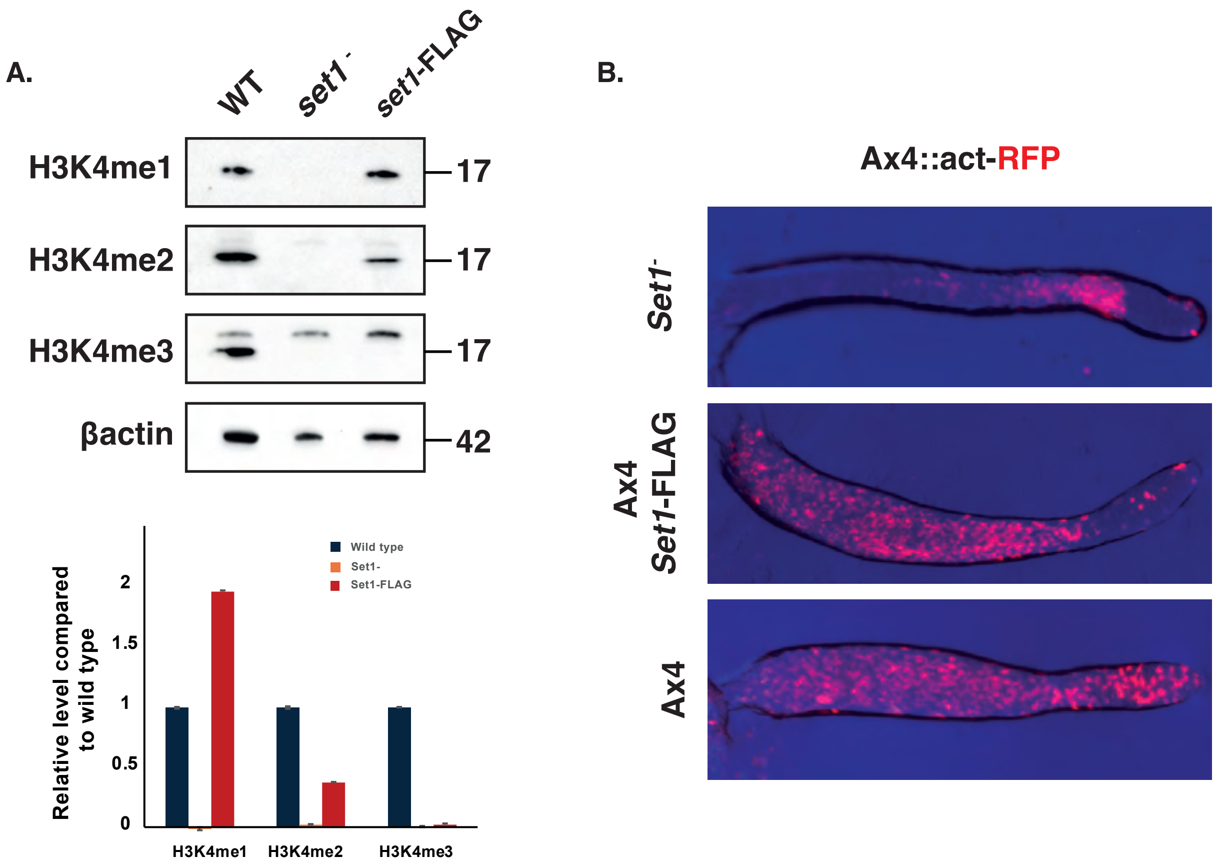

#### Supplementary figure 10

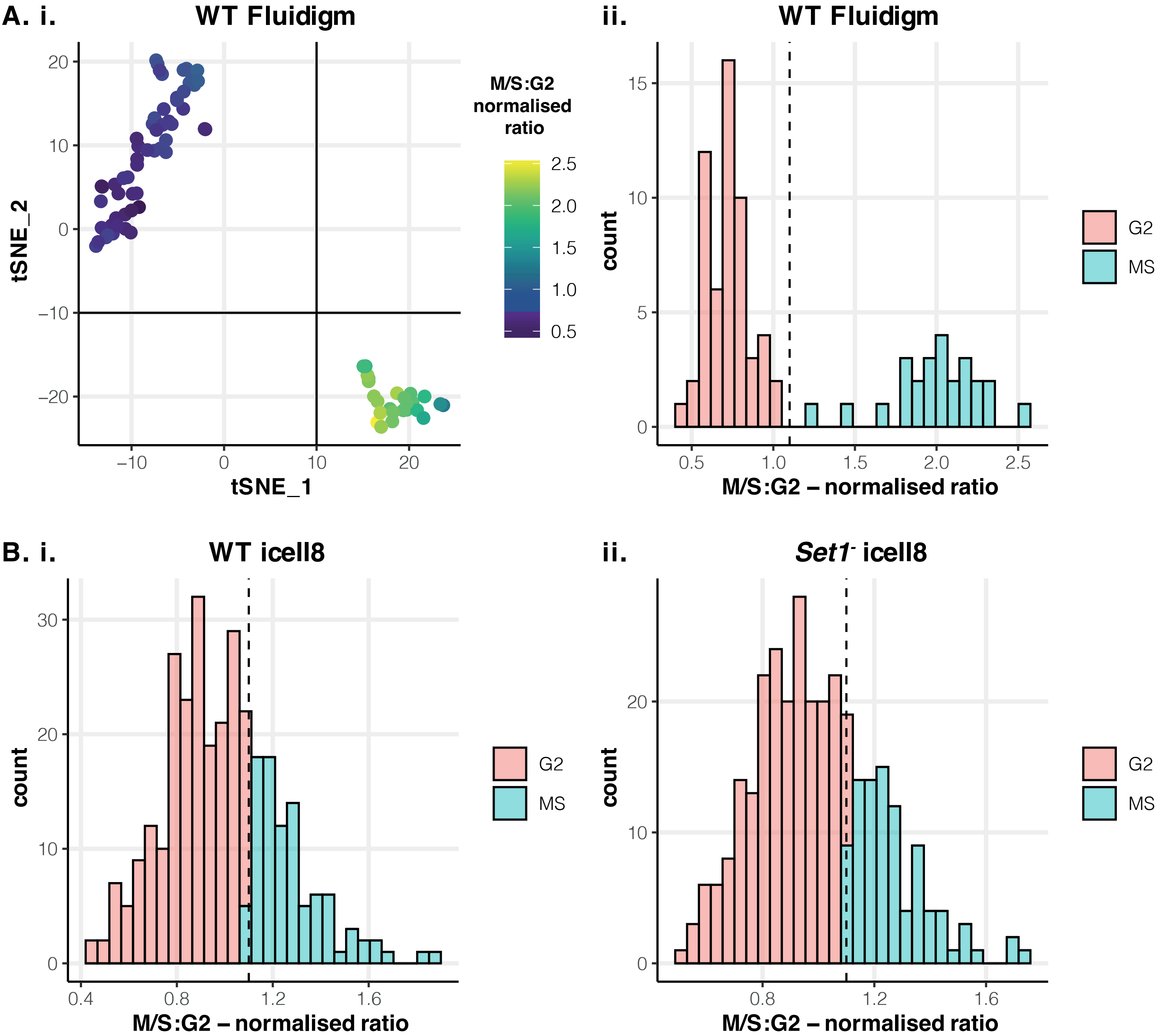

#### Supplementary figure 11

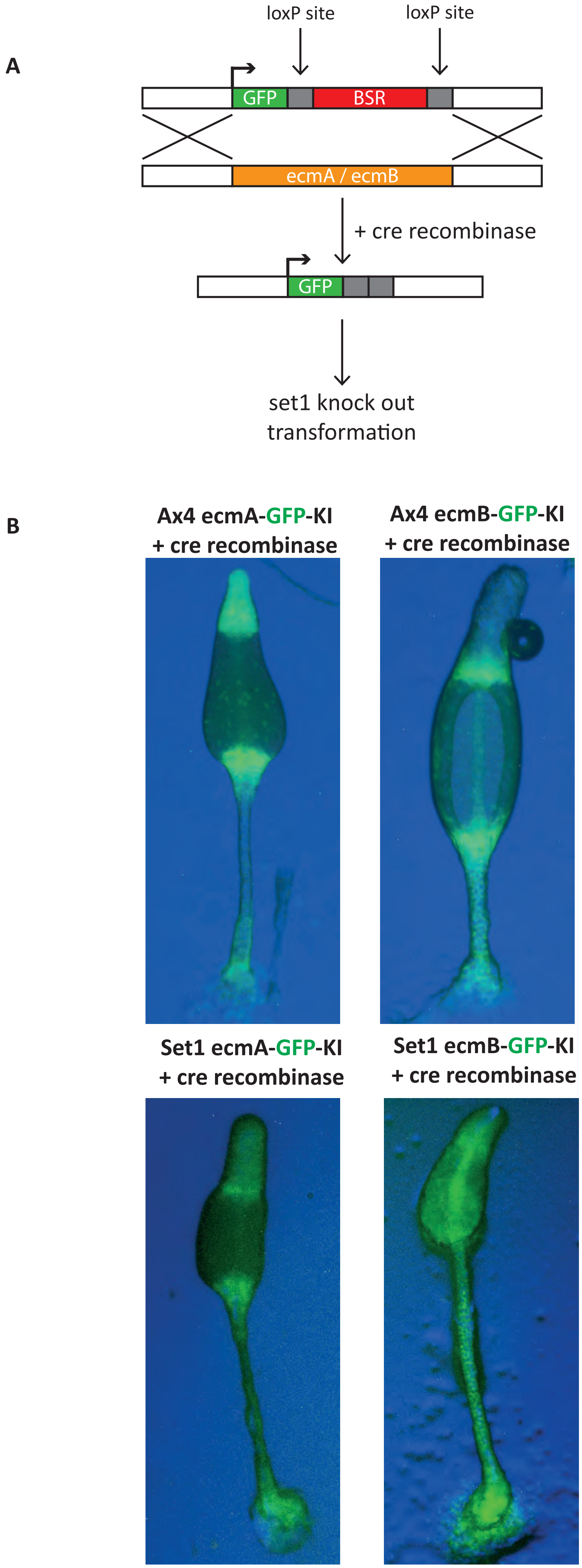

Supplementary figure 12

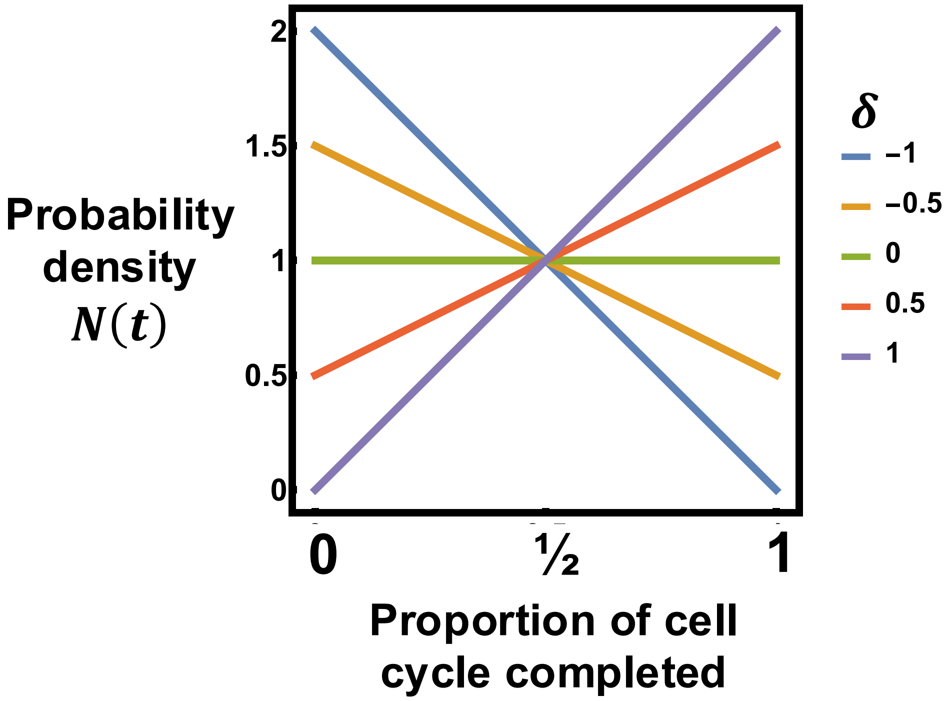

Supplementary figure 13

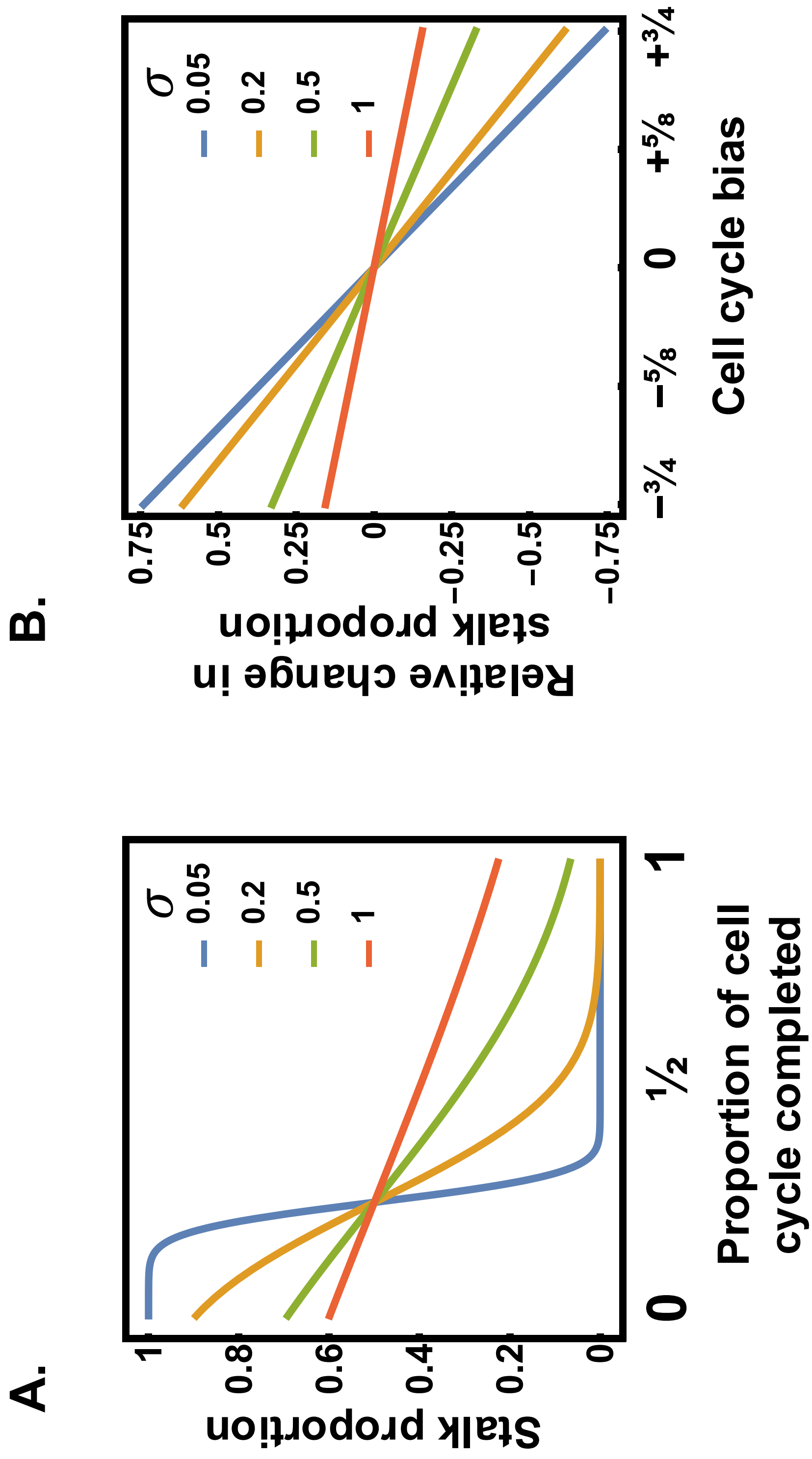
